# Diabetic Retinopathy detection through integration of Deep Learning classification framework

**DOI:** 10.1101/225508

**Authors:** Alexander Rakhlin

**Affiliations:** Neuromation OU Tallinn, 10111 Estonia

**Keywords:** Diabetic Retinopathy, Deep Learning, Medical Imaging, Computer-aided diagnosis (CAD), Image Recognition

## Abstract

This document represents a brief account of ongoing project for Diabetic Retinopathy Detection (DRD) through integration of state-of the art Deep Learning methods. We make use of deep Convolutional Neural Networks (CNNs), which have proven revolutionary in multiple fields of computer vision including medical imaging, and we bring their power to the diagnosis of eye fundus images. For training our models we used publicly available Kaggle data set. For testing we used portion of Kaggle data withheld from training and Messidor-2 reference standard. Neither withheld Kaggle images, nor Messidor-2 were used for training. For Messidor-2 we achieved sensitivity 99%, specificity 71%, and AUC 0.97. These results close to recent state-of-the-art models trained on much larger data sets and surpass average results of diabetic retinopathy screening when performed by trained optometrists. With continuous development of our Deep Learning models we expect to further increase the accuracy of the method and expand it to cataract and glaucoma diagnostics.

## 1 Introduction

Diabetic Retinopathy (DR) is the leading cause of blindness in the working-age group. Among 23 million Americans, 59 million Europeans, and as many as 50 million Indians suffering from diabetes, the prevalence of those with DR is estimated between 18% and 28%. Regular eye examination among these vulnerable groups is necessary to diagnose DR at an early stage, when it can be treated with the best prognosis. Currently, detecting DR is a time-consuming and manual process that requires a trained clinician to examine and evaluate digital color fundus photographs of the retina. The clinical grading process consists of detection certain subtle features, such as microaneurysms, exudates, intra-retinal hemorrhages and sometimes their position relative to each other on images of the eye.

**Figure 1:**
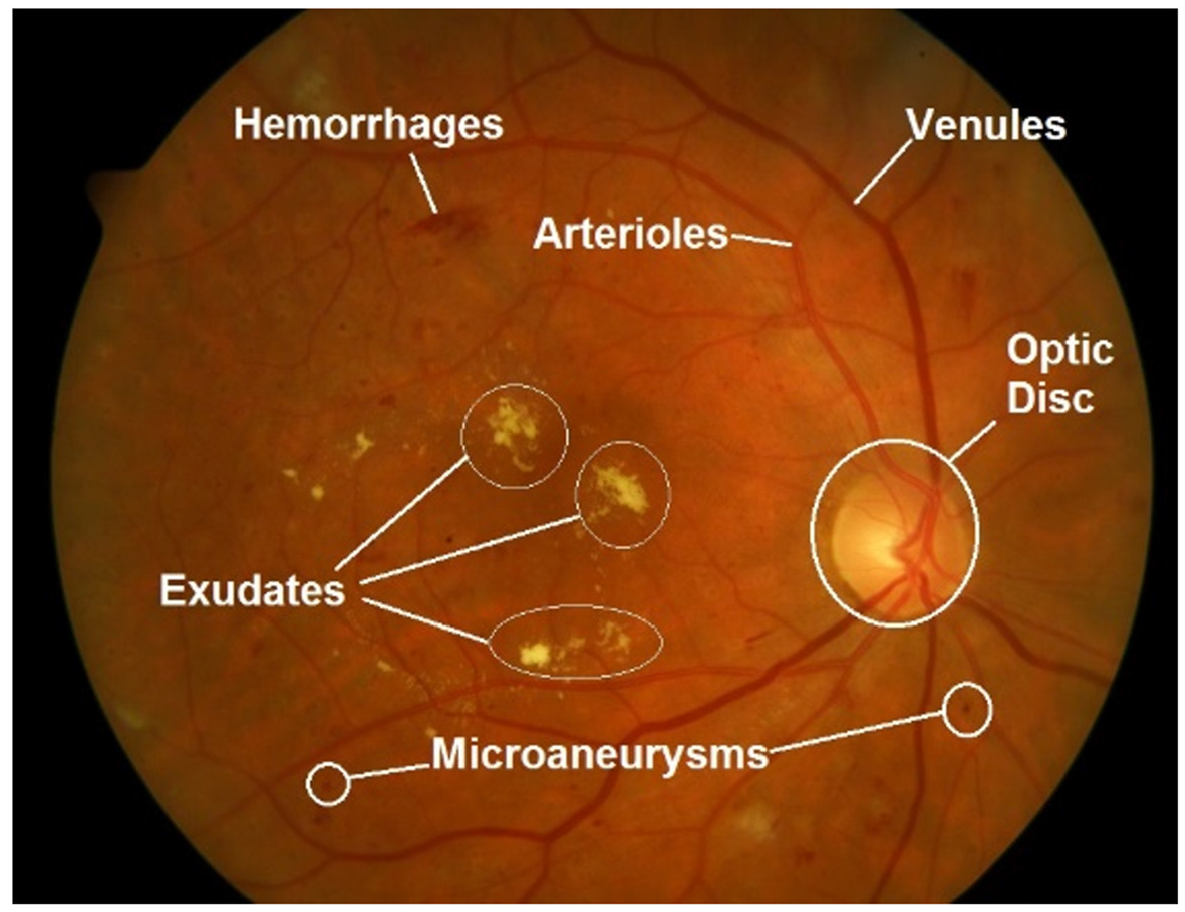
Abdullah et al. (2016)

Automated analysis of retinal color images has such benefits as increased efficiency and coverage of screening programs, reduced barriers to access, and early detection and treatment. Over the last several decades the algorithms for automated retinal screening have been developed. Until recently, those algorithms have been using analysis based on image features manually crafted by experts. Deep Learning is an approach that avoids such engineering automatically learning hierarchy of discriminative features directly from the images given a large set of labelled examples. With AlexNet [1] stealing the show in 2012 Deep Convolutional Neural Network (CNN) have revolutionized the field of computer vision and have been highly successful in a large number of computer vision and image analysis tasks, substantially outperforming all classical image analysis techniques. In the domain of retinal image analysis, CNNs have been used for vessel segmentation to classify patch features into different vessel classes [5]. In the Kaggle competition [4] all top solutions used CNNs to identify signs of DR in retinal images.

In this project we use Deep Learning models to detect referable Diabetic Retinopathy (rDR) in 2 data sets. We assess the sensitivity, specificity and area under the operator receiving characteristics curve (AUC) of the model, and compare these with other state-of-the-art models and clinicians.

## 2 Data

For the present project we used two data sets of high-resolution retinal color images.

**Kaggle data set** [4] was provided by EyePACS, a free platform for retinopathy screening. The data set consists of 88,696 images of 44,348 subjects, one image for each eye. The images in this dataset come from different models and types of cameras and feature very mixed quality. A clinician has rated the presence of diabetic retinopathy in each image on a scale of 0 to 4, according to International Clinical Diabetic Retinopathy severity scale (ICDR):

0 – No DR
1 – Mild DR
2 – Moderate DR
3 – Severe DR
4 – Proliferative DR

We consider a patient as referable if the DR stage is between 1 and 4, otherwise we consider the patient as non-referable. We created two levels of disease for each image:

0 – No DR – ICDR level 0
1 – Referable DR (rDR) – ICDR level 1 to 4

Kaggle image quality was estimated as 75% gradable. rDR prevalence in Kaggle data set was 30.5% (13,545 subjects, 27,090 images). Kaggle data set was randomly split into two uneven subsets. 81,670 retinal images of 40,835 subjects were used for model development. 7,026 images of the other 3,513 subjects were used for model evaluation.

**Messidor-2** [3] is publicly available dataset which has been used by other groups for benchmarking performance of algorithms for DRD. The corresponding rDR reference standard is available for researchers [6]. Messidor-2 consists of 1,748 retinal color images of 874 subjects. Subjects were imaged using a color video 3CCD camera on a Topcon TRC NW6 non-mydriatic fundus camera with a 45 degree field of view, centered on the fovea, at 1440×960, 2240×1488 or 2304×1536 pixels. Messidor-2 was 100% gradable. rDR prevalence was 21.7% (190 subjects, 380 images). Entire Messidor-2 was used for model evaluation.

**Notice of image quality.** Retinal images are acquired using different hardware operated by specialists who have varying levels of experience. This results in a large variation in image quality. Subtle signs of retinopathy at an early stage can be easily masked on a low contrast, or blurred, or low resolution image (see Fig. 2). Analysis of an image of low quality may produce unreliable results when the system labels an image as normal while lesions are present. That is why image quality is very important factor for DRD system sensitivity.

**Figure 2:**
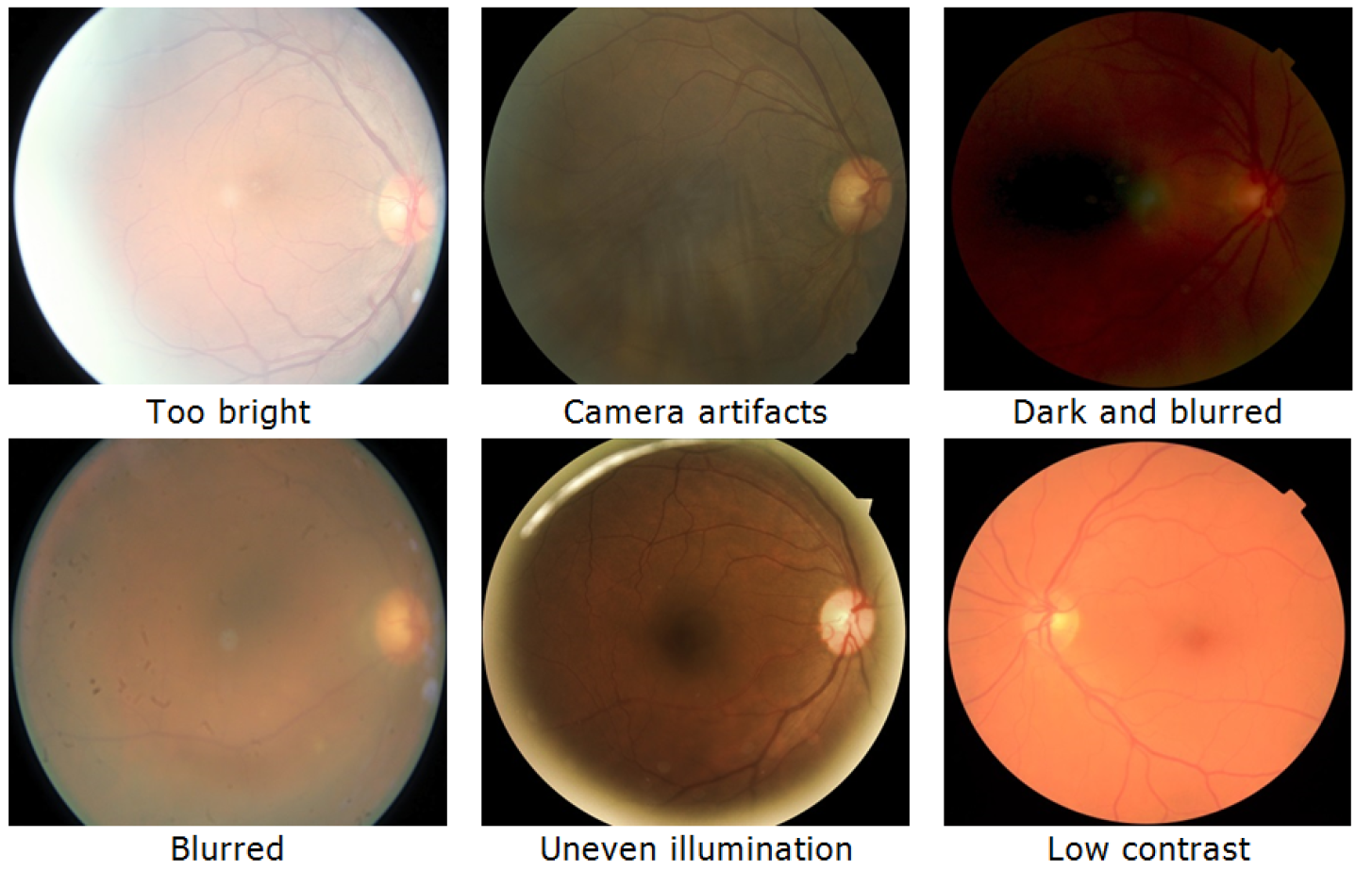
Retinal images of deficient quality

## 3 Method

There are three major tasks in computer vision in increasing order of difficulty: (I) classification, (II) localization/segmentation, and (III) detection. *Detection task* represents the highest difficulty and requires detection of many small objects, while *localization task* pertains to a single, usually large object. In *classification task* an image is assigned a single label or score corresponding to the image as a whole, Fig. 3.

**Figure 3:**
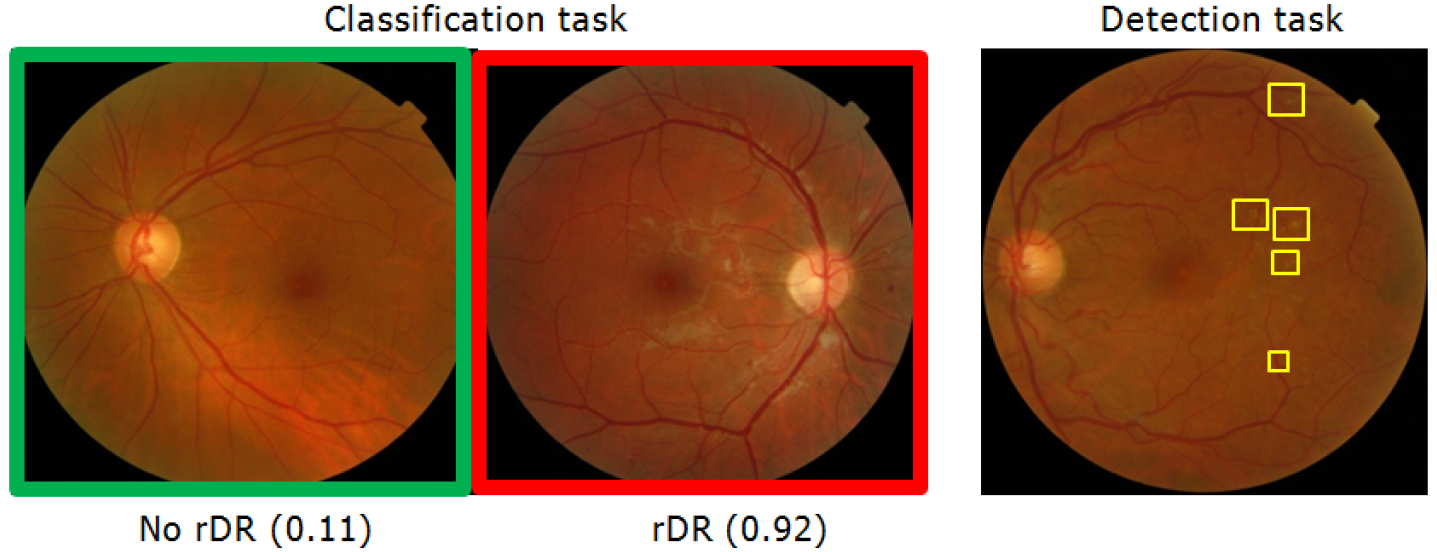
Classification vs. Detection

In DRD project, advantage of the classification task is that it does not require
manual annotation of retina regions (as opposed to detection task), binary or categorical labels is enough for training a model. However, one serious challenge for classification task comes from the fact that on global scale healthy retinal images do not differentiate from those with DR. It is subtle lesions (Drusen, Exudates, Microaneurysms, Hemorrhages, and Cotton-wool Spots) observed on image patch scale make the difference, Fig. 4.

**Figure 4:**
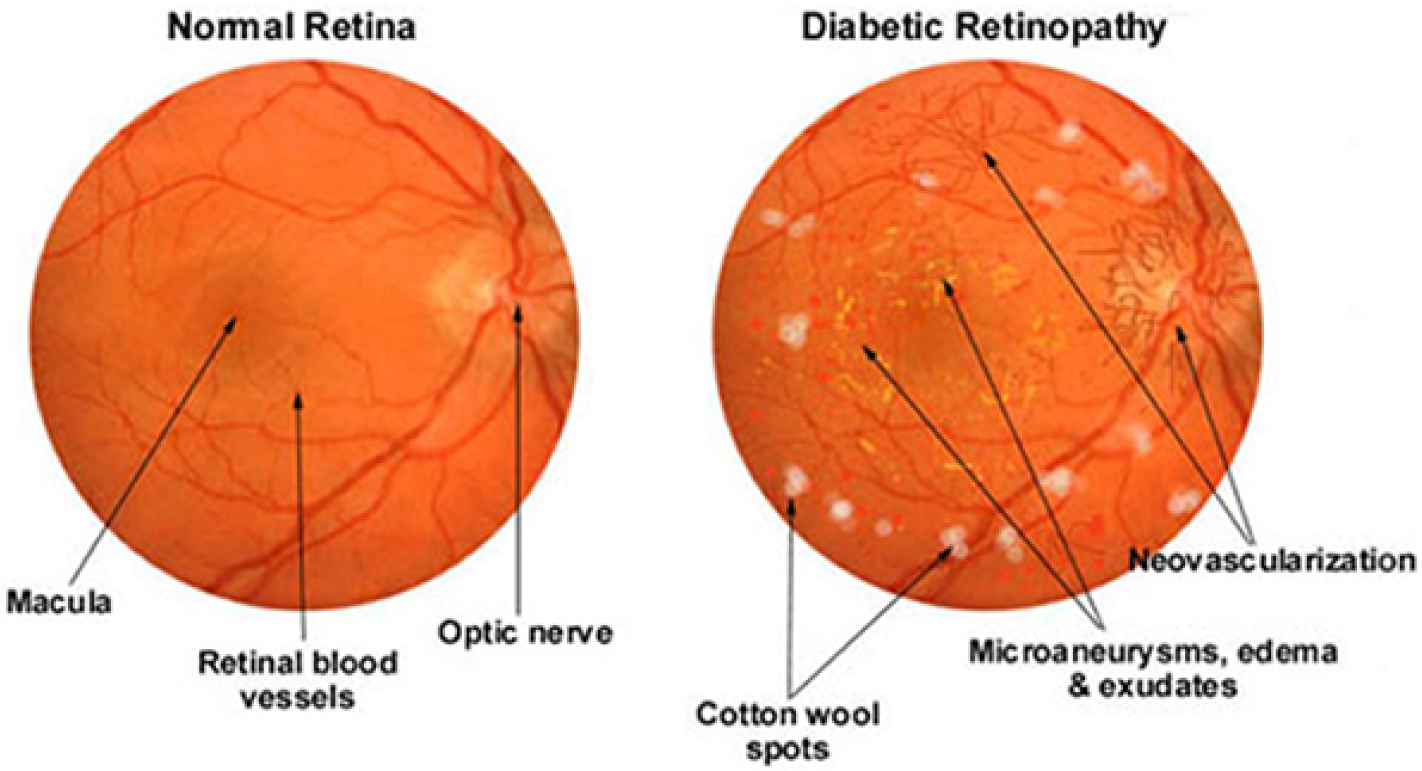
Microaneurysms, Exudates and Cotton-wool Spots source by: http://www.coatswortheyeclinic.co.uk/photography/3102576

Fortunately, with recent advance of Deep and extra-Deep Convolutional Neural Networks (CNNs), it became possible to train powerful classification models capable of automatic discovery subtle local features without need of manual annotation of individual lesions. The network used in our project is a Convolutional Neural Network with deep layered structure that combines nearby pixels into local features, and then progressively aggregates those into global hierarchical features. The network was trained on binary classification task and outputs continuous score between 0 and 1, which represents classifier’s confidence in Referable DR presence. Although classification model does not explicitly detect lesions (Drusen, Exudates, Microaneurisms, Hemorrhages, or Cotton-wool Spots), it likely learns to recognize them using the local features. The architecture used in our project originates from famous VGGNet family, winner of ImageNet Challenge ILSVRC-2014, designed for large-scale natural image classification [2], Fig. 5.

Training convolutional model 19 layers deep, with 8,013,393 parameters, on large 540×540 images represents significant challenge for convergence. We significantly redesigned original VGG architecture, added number of recent innovations and developed special training protocols to achieve better convergence and accuracy. More traditional techniques like dropout, data augmentation and preprocessing have been applied too. With these design choices, full end-to-end training is possible, and for a single model it takes several days on Kaggle data set on GPU Nvidia GTX-980.

**Figure 5:**
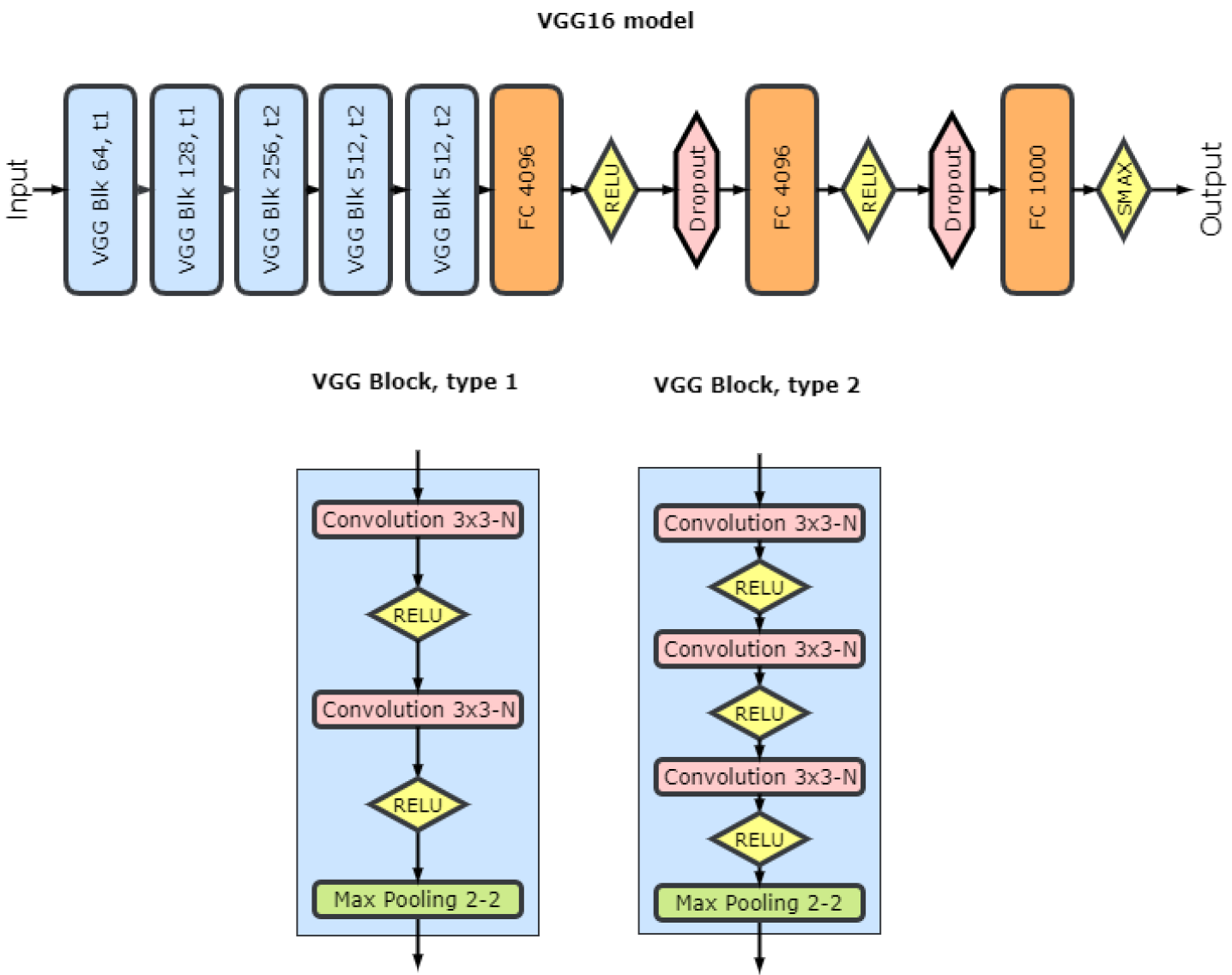
Original VGG16 model

### 3.1 Diagnostic pipeline

Unlike training, deployment and application of the model is straightforward, fast and not resource intensive. In principle, it can run on CPU, though GPU is recommended.

1. First, we preprocess retinal images to make them uniform. We normalize [^1^ We consider a number of normalization techniques like color, contrast normalization,whitening (sphering)], scale, center and crop them to 540×540 pixels.
2. Next stage is image quality assessment module [^2^ In development]. For DR detection system sensitivity is a key factor. Subtle signs of retinopathy at an early stage can be easily masked on a low contrast or blurred image. Analysis of an image of low quality may produce unreliable results when the system labels an image as normal while lesions are present. These low quality images should be automatically detected, and examined by a specialist, and reacquired if necessary.
3. Then we generate a number of randomly augmented images and feed them into DRD model(s).
4. Multiple scores obtained on previous stage are being fused into final diagnosis.
5. When possible, we combine the other eye scores too – for added accuracy.

**Figure 6:**
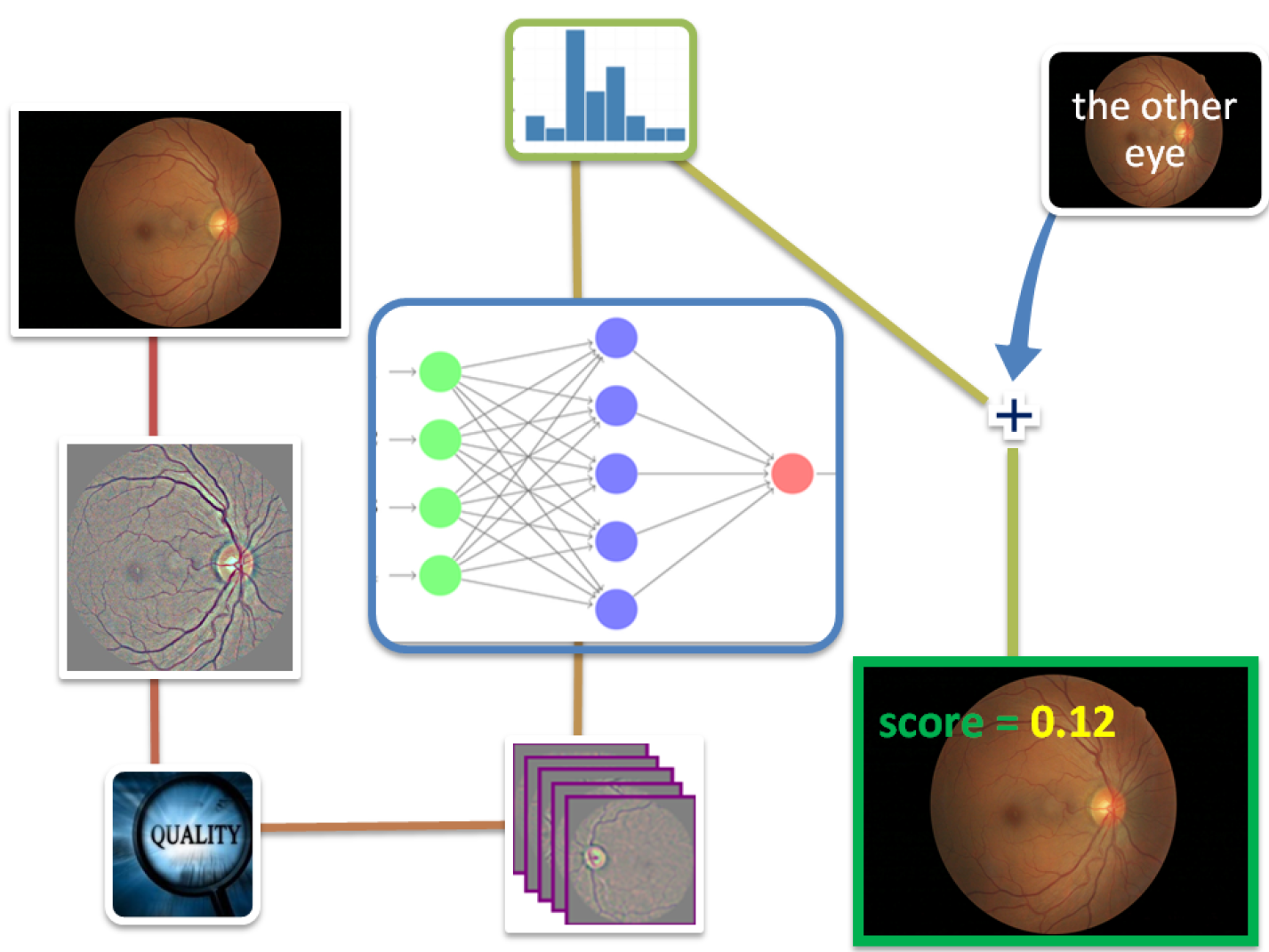
Diagnostic pipeline

### 3.2 Limitations and applicability

Just like any automated system, even powerful Machine Learning models have their theoretical limits. Apart from specific quality considerations we discussed above, they demonstrate best performance being applied to images of the same *genesis* as those they were trained on. This means that images obtained using very different hardware, or from peculiar population not present on training stage, may sometimes result in decreased accuracy. Fortunately, these limitations can be circumvented by means of fine-tuning the model on the new data, or via training on broader data sets.

## 4 Results and conclusions

We tested the model on 2 data sets, Messidor-2 and portion of Kaggle withheld from training. The model was never trained on any of the images from these data sets. Metz ROC Software package [7] was used to estimate area under the ROC curve (AUC) of the model for detecting rDR and its Confidence Intervals. Two operating points have been selected, one for high sensitivity and another for high specificity. Fig. 7

**Figure 7:**
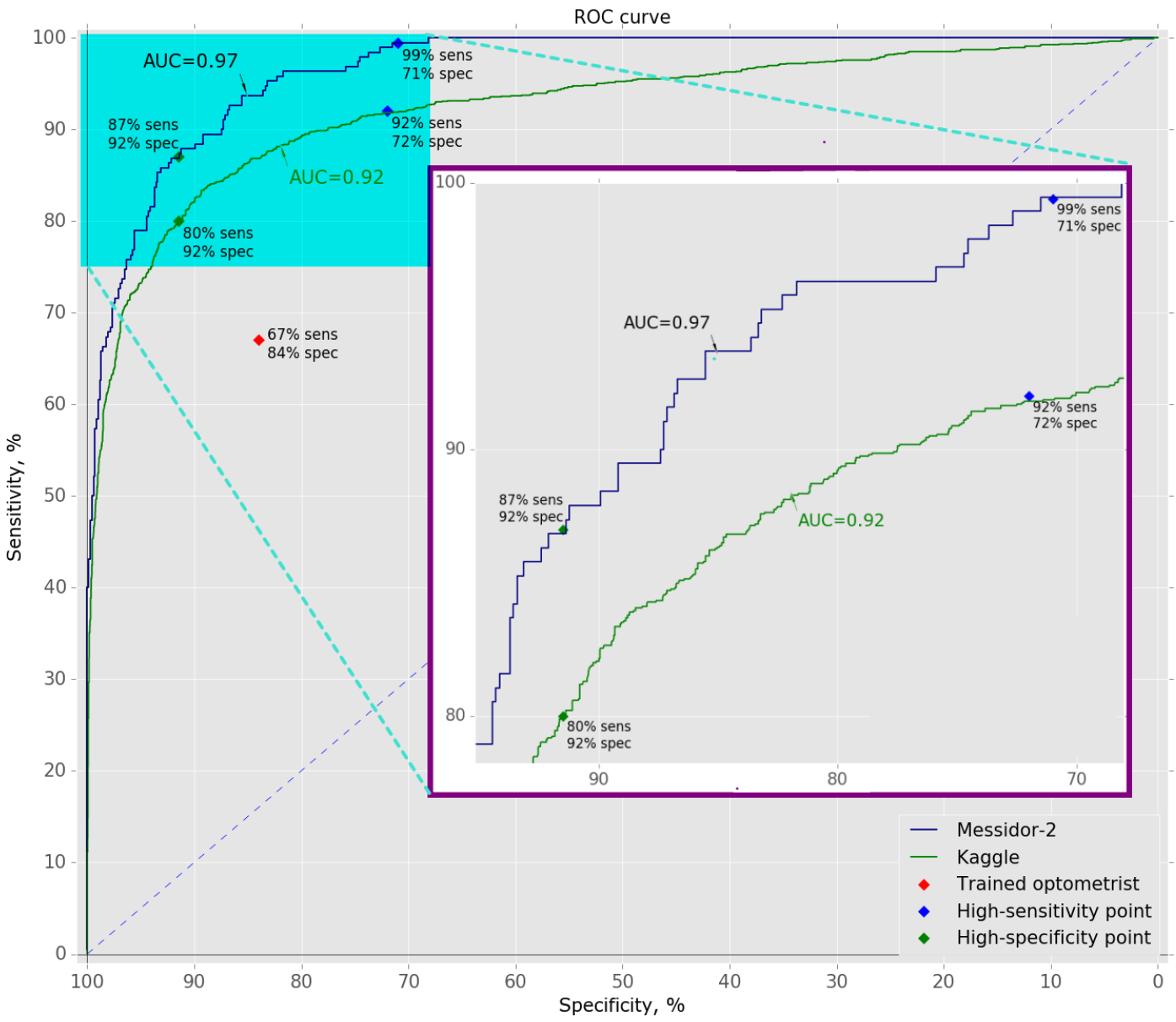
Receiver Operating Characteristic of the Model

**Messidor-2.** For 874 subjects, the area under the ROC curve was 0.967 (95% CI: 0.959 - 0.974). The sensitivity/specificity pair of the model was 99/71% at high sensitivity operating point. At high specificity operating point sensitivity/specificity was 87/92%

**Kaggle.** For 3,513 subjects, the AUC was 0.923 (95% CI: 0.915- 0.931). The sensitivity/specificity of the model was 92/72% at high sensitivity operating point. At high specificity operating point sensitivity/specificity was 80/92% (table 1)

These results close to recent state-of-the-art models [8, 9], trained on much larger data sets and surpass average results of diabetic retinopathy screening when performed by trained optometrists [10]. One interesting observation is different definitions of Referable DR we use. Our definition of rDR is ICDR level 1 to 4. The state-of-the-art models define rDR as ICDR level 2 to 4. This might suggest that our model is more sensitive to DR at an early stage; however we have not had an opportunity to verify this hypothesis.

**Table 1.** Results

| Data set | AUC | High sensitivity OP |  | High specificity OP |  |
| --- | --- | --- | --- | --- | --- |
|  |  | Sensitivity | Specificity | Sensitivity | Specificity |
| Messidor-2 | 0.97 | 99% | 71% | 87% | 92% |
| Kaggle | 0.92 | 92% | 72% | 80% | 92% |

Why the model performance on Messidor-2 is superior to Kaggle? Messidor data set used in this project consists of 100% gradable retinal images of high quality. Kaggle’s quality is very mixed and estimated as 75% gradable. This is the reason why image quality assessment module is of high importance and priority. Thereby, Kaggle is more representative of data from screening programs than Messidor-2 is. In the same time, Messidor-2 is a “golden standard” as it has been used by other groups for benchmarking performance of automated detection algorithms for diabetic retinopathy.

Finally, are our operating points (cut-offs) suitable in any scenario? Not likely. Depending on the prevalence of rDR in the population and medical objectives, another trade-off between sensitivity and specificity could be found more appropriate.

## 5 Future Plans

The project is constantly evolving. Implementing novel techniques, we expect to further increase the accuracy of the method as well as to expand it to other eye diseases. Here is an abbreviated list of steps we take to achieve our objectives:

- larger data sets for model development
- image quality assessment module
- experiments with other deep learning architectures
- development of a neural model based on detection principles and capable of exact discovery of subtle lesions (Drusen, Exudates, Microaneurysms, Hemorrhages, and Cotton-wool Spots) just like retinal specialists do
- expansion to cataract and glaucoma diagnostics

## Appendix A

Here we provide some technical details of the implementation. The project is being developed in Python 3.4 environment and makes use of the following libraries, platforms and hardware:

- Keras v1.2, Deep Learning library for Theano and TensorFlow
- Theano v0.8, a Python framework for fast computing
- cv2 v3.1, a Python wrapper for OpenCV
- SciPy v0.18, a Python-based ecosystem of open-source software for mathematics, science, and engineering.
- NumPy v1.11, the fundamental package for scientific computing with Python
- Pandas v0.18, a data analysis and manipulation library for Python
- Matplotlib v1.5, a Python plotting library
- CUDA v7.5, a GPU computing platform
- CuDNN v5.0, the NVIDIA GPU-accelerated library
- Intel Core i5-6500 @3.2 GHz, 16GB RAM, NVIDIA GeForce GTX 980 Graphic Card

